# Scanning for Representation: A Scoping Review of Racial and Ethnic Diversity in MRI Studies of the Maternal Brain

**DOI:** 10.1101/2025.06.20.660240

**Authors:** Edwina R. Orchard, Clíona L. Murray, Kathryn M. Wall, Josephine C.P. Levy, Jin Young Shin, Claudia G. Gaebler, Victoria R. Hart-Derrick, Kathy Ayala, Amorine Adodo, Michèle J. Day, Melissa C. Funaro, Kathleen Guan, Termara C. Parker, Jocelyn A. Ricard, Francesca Penner, Tal Yatziv, Helena J.V. Rutherford

## Abstract

A growing number of magnetic resonance imaging (MRI) studies are examining brain changes across pregnancy and early motherhood, gaining fundamental insight into the neural adaptations of motherhood, with critical clinical and policy implications for supporting mother, child, and family unit. As the field takes off, now is the time to take stock of the current literature and neuroscience practices, to ensure that the field is based on studies that are robust, representative, and transparent. Here, we conducted a scoping review to understand the racial and ethnic diversity of participants reported in MRI studies of the maternal brain, guided by the Joanna Briggs Institute methodology. Our findings highlight three key issues in the 185 identified studies of the maternal brain using MRI: (1) the widespread underreporting of participant racial and ethnic data, with only 38.38% of studies reporting race and/or ethnicity demographics; (2) the overrepresentation of white participants, with 46.83% of the samples that report race and/or ethnicity identifying as white/Caucasian; and (3) the disproportionate geographical locations of studies, with 68.65% of studies from North America or Europe and Central Asia. These findings raise concerns about the generalizability of existing research beyond WEIRD (western, educated, industrialized, rich and democratic) populations, and underscore the urgent need for concerted structural change in neuroscience research practices. While identifying a lack of diversity is only the first step, this scoping review serves as a call to action for greater representation in future research, for our own research group as well as others.

## Introduction

The neuroscience of human motherhood is a field in its infancy, with a growing number of studies using neuroimaging to examine brain changes across pregnancy and early motherhood. These studies suggest that structural and functional brain changes support sensitive maternal caregiving behaviors^1–3^, for example, heightened neural responses to audio/visual infant cues, which may be altered in mothers with postpartum psychopathology^4–8^. Increasing our understanding of the maternal brain has the potential to improve maternal health and wellbeing, with long-lasting^9^ positive impacts for mother, child, and family unit. However, the benefit and generalizability of the maternal brain literature may be limited by current and historic systematic racial bias and exclusionary practices in neuroscience.

Neuroscience research has historically excluded minoritized people^10–14^. As such, our understanding of the human brain is disproportionately based on samples from Western, educated, industrialized, rich, and democratic (WEIRD) populations^15,16^, with an insufficient representation of racially and/or ethnically minoritized people. Such biased sampling distorts our understanding of the brain, and renders research findings less accurate, generalizable, and reproducible^17,18^, making it not only misguided, but actively harmful to assume that research findings will generalize beyond these WEIRD samples. We cannot use neuroscience findings to inform equitable clinical practice and public policy if there are disparities in the utility and benefits of research findings for minoritized populations^16,18^.

It is crucial to understand that race is a socially-determined construct^19,20^, with racial boundaries historically prescribed to reflect economic, cultural, political, and social structures, that vary across regions, cultures, or periods in time, and uphold systems of colonialism and imperialism^16^. However, despite the social construction of race and ethnicity, tangible racial disparities in health outcomes continue to exist, including the racial disparities related to maternal health and wellbeing. For example, in the United States, Indigenous women are twice as likely, and Black women are three times more likely, to die from a pregnancy-related cause compared to white women^21–23^. These fatal disparities persist beyond differences in income and education^23^. These health inequalities are beginning to be understood as consequences of the *experience* of systemic racism and structural inequities, not race itself^11^, highlighting the importance of dismantling systems of oppression and marginalization in healthcare, as well as in research.

Magnetic resonance imaging (MRI) is increasingly used to examine the structure and function of the maternal brain^24^, gaining considerable public interest in response to high-impact publications and science communication efforts in popular-science books^25–27^, podcasts^28,29^, and mainstream media coverage^30^. Given the potential for this field to impact public policy, medical practice, and public opinion, it is crucial that equipment is inclusive, samples are diverse, and research findings are representative. A recent review of racial and ethnic representation in electroencephalography (EEG) studies of the maternal brain found a disproportional over-representation of participants that identified as white/of European ancestry, and were mostly conducted in the United States and Europe^31^. This suggests that the current EEG literature on the maternal brain may not generalize across racially diverse populations. Importantly, EEG has been heavily critiqued beyond the maternal brain literature for exclusionary methodology, restricted by hair texture and density, such as coarse and curly hair types common in individuals of African descent, leading to the disproportionate exclusion of Black participants^10,12,17,32^. Similarly, pulse oximeters and functional near-infrared spectroscopy (fNIRS) have been shown to be less effective for darker and more pigmented skin complexions^14,33–35^. MRI has been similarly critiqued for methodological limitations which exclude racially and ethnically diverse populations. For example, the MRI head coil can restrict the ability to accommodate natural hairstyles (i.e., afro-textured hair or braids). Additionally, metallic objects pose a safety risk, and result in artifacts for MRI scans^36^, including the metal found in hairstyles such as weaves, sew-in hair extensions, and/or braids, which are commonly worn by Black women^12^, and in the pins and clips used to secure hijabs and other head coverings. Issues of diversity and representation are exacerbated when researchers do not prioritise culturally-inclusive approaches to study design, participant recruitment, data collection, analysis, and interpretation^13^.

In this scoping review, we examine the reporting of racial and/or ethnic information in studies of the maternal brain using MRI in pregnancy and the postpartum period. Given the limited diversity of maternal EEG studies^31^, we hypothesize that there will also be limited racial and/or ethnic diversity in studies of the maternal brain using MRI. We analyze and discuss these findings in the context of the geographic locations where the studies were completed, to reflect that: (1) the racial and/or ethnic compositions of different populations vary considerably due to ancestral origins, voluntary and forced migration, colonization, enslavement, trade, conflict, environmental shifts, and modern globalization; (2) as race and ethnicity are socially-determined, the meaning and conceptualization of these constructs vary geographically and intersect with language, culture and intersectional identity; and (3) there are important differences in social norms and legal restrictions surrounding the collection and reporting of race and/or ethnicity data in different countries and regions.

## Methods

This scoping review was conducted in accordance with the Joanna Briggs Institute (JBI) methodology for scoping reviews^37^ and the Preferred Reporting Items for Systematic Reviews and Meta-Analysis Extension for Scoping Reviews (PRISMA-ScR) Checklist and Explanation document^38^. The review protocol was registered with the Open Science Framework on December 19th, 2022 (https://osf.io/6c7zt).

### Review Questions

This scoping review was guided by several research questions: What proportion of MRI studies of the maternal brain collect and report the racial and ethnic demographics of their sample? What are the racial and ethnic compositions of these MRI studies of the maternal brain? What are the geographic locations of MRI studies of the maternal brain? Do the racial and ethnic compositions of MRI studies of the maternal brain represent the composition found in the general population(s) where the data were collected? What proportion of MRI studies of the maternal brain incorporate racial and ethnic demographics into their statistical analyses?

### Eligibility

The inclusion criteria, defined in accordance with the JBI recommendations for scoping reviews, were as follows:

1. Participants: Human mothers, who were assessed during pregnancy and/or in the postpartum period.
2. Concept: Examination of studies which employed structural and/or functional magnetic resonance imaging (MRI) to investigate the maternal brain.
3. Context: Experiments conducted in any setting, independent of the country of the study.
4. Type of Sources: Peer reviewed published papers utilizing quantitative and mixed methods study designs, including studies which use between-groups comparisons to examine differences between mothers and non-mothers, pregnant and non-pregnant women, or between mothers and fathers, as well as within-groups designs examining mothers longitudinally, and studies which examine associations between the maternal brain and other maternal factors (e.g., psychopathology or maternal caregiving behaviors).

The exclusion criteria were as follows:

1. Non-empirical research papers including reviews, book chapters, and commentaries/opinion pieces.
2. Studies which used neuroimaging modalities other than MRI (e.g., Positron Emission Tomography [PET], EEG, fNIRS).
3. Studies which did not scan mothers’ brains using MRI (e.g., assessment of non-mother women only, inclusion of fathers only, mothers included in study but whose brains were not scanned using MRI).
4. Studies in which participants were limited to mothers beyond the postpartum period. The end of the postpartum period was defined for this purpose as the mean of the sample having a youngest child under 3 years (36 months) of age.
5. Case studies and medical studies which examined medically indicated MRI scans only (e.g., in the case of stroke).
6. Non-peer reviewed/non-published papers (e.g., popular-science books, theses/dissertations, protocol papers, preprints, retracted papers, amendments/ corrections/corrigenda, conference abstracts).
7. Papers not available in the English language.

### Search Strategy

To identify relevant literature, the following databases were searched: MEDLINE, Embase, PsycInfo, Web of Science, and Cochrane with the help of an experienced librarian (author M.C.F.). The systematic search was conducted for the period from database establishment to November 11, 2022, with no language limitations. The search was rerun on October 12, 2023, to capture additional literature published during the review process. A medical subject heading (MeSH) analysis of known key articles on maternal brain MRI studies was conducted. An iterative process was used to translate and refine search terms. To maximize sensitivity, the formal search used controlled vocabulary terms and synonymous free-text words. The search strategy was peer reviewed by a second librarian, not associated with the project, using the PRESS standard^39^. The comprehensive search strategies for all databases are included in Supplementary Table 1. Reviewers checked the included studies for additional relevant cited and citing articles using the citationchaser software^40^.

### Study Selection

Upon search completion, all identified citations were collated and uploaded into *EndNote* (v20.4), and duplicates were removed using the Yale Reference Deduplicator (https://library.medicine.yale.edu/reference-deduplicator) and uploaded into Covidence (https://www.covidence.org/) for screening. Titles and abstracts were independently screened by two or more independent reviewers for assessment against the inclusion criteria for the review. The full text of selected citations was then assessed in detail against the inclusion criteria by two independent reviewers. Any disagreements that arose between the reviewers at each stage of the selection process were resolved through group discussion or an additional reviewer. This process resulted in 185 articles meeting inclusion criteria (Figure 1).

**Figure 1:**
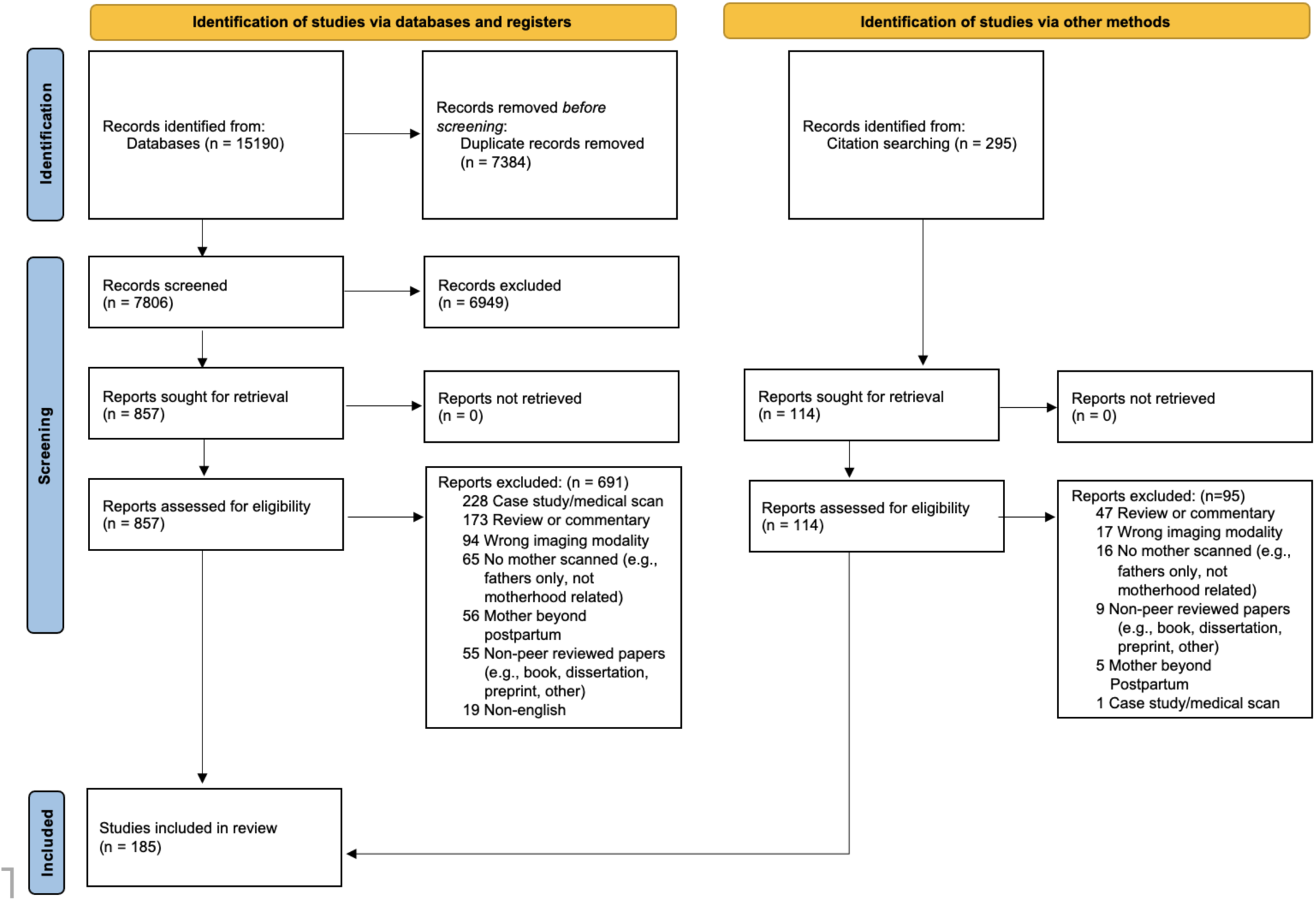
Preferred Reporting Items for Systematic Reviews and Meta-Analysis Extension for Scoping Reviews. (PRISMA-ScR) flow diagram of study selection resulting in 185 included papers.

### Data Extraction

Data were extracted from articles included in the scoping review by one independent reviewer using a data extraction tool developed by the reviewers, and each extraction was checked for consistency and errors by author E.R.O. Some studies reported the demographics of their sample at the total participant level, reporting race and/or ethnicity numbers or percentages that were not distinguishable between groups (e.g., reporting mothers’ and non-mothers’ demographics together). For this reason, we chose to include data for all participants in the identified studies, reflecting the influence and importance of also having representative control/comparison groups on our understanding of the maternal brain.

As the majority of studies that reported race and/or ethnicity were from the United States (53/ 71 studies; 74.65%), racial and ethnic data were extracted verbatim, and then collated into categories aligned with the United States’ census reporting guidelines. These included American Indian/Alaskan Native, Asian, Black/African-American, Pacific Islander/Native Hawai’ian, white/Caucasian, Two or More Races/Ethnicities. In the included studies, there was inconsistent reporting of Hispanic/Latine identities (i.e., sometimes reported as ‘race,’ and sometimes as ‘ethnicity’). In the current review, we have reported Hispanic/Latine data both within the racial demographics, and as distinct ethnicity demographics, to accurately report the available data. In addition to these census categories, we also created three more categories: ‘Eurocentric’, ‘Other’ and ‘Not Reported/Unspecified’. The Eurocentric label was created to describe participants labelled relative to whiteness, including participants “of minority race” or “non-white” participants. Table 1 details the verbatim phrasing used within papers, and our coding scheme used to aggregate reported demographics into consistent race/ethnicity variables.

**Table 1:**
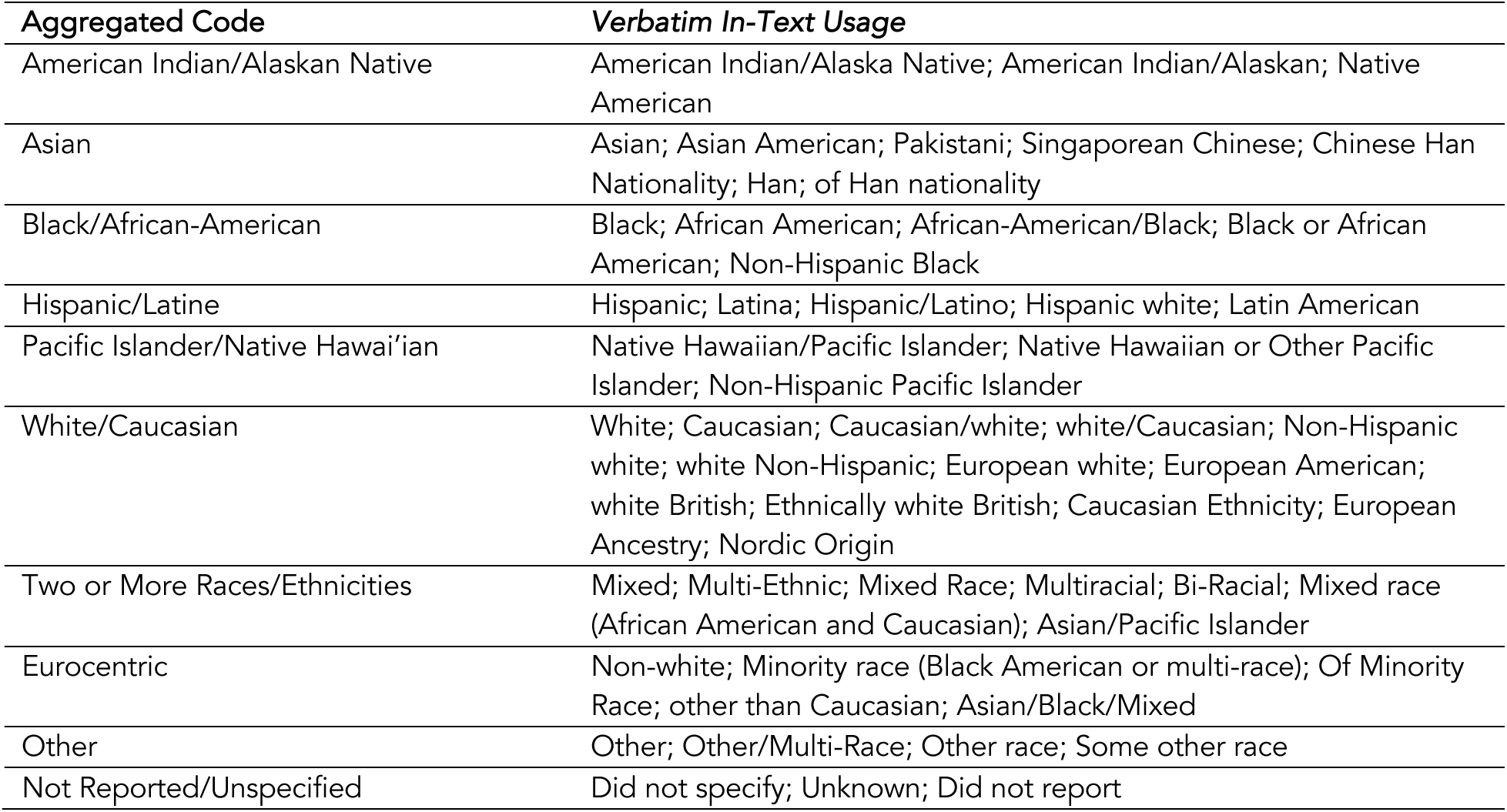
Coding scheme for aggregated race and/or ethnicity variables.

| **Aggregated Code** | **Verbatim In-Text Usage** |
| --- | --- |
| American Indian/Alaskan Native | American Indian/Alaska Native; American Indian/Alaskan; Native American |
| Asian | Asian; Asian American; Pakistani; Singaporean Chinese; Chinese Han Nationality; Han; of Han nationality |
| Black/African-American | Black; African American; African-American/Black; Black or African American; Non-Hispanic Black |
| Hispanic/Latine | Hispanic; Latina; Hispanic/Latino; Hispanic white; Latin American |
| Pacific Islander/Native Hawai'ian | Native Hawaiian/Pacific Islander; Native Hawaiian or Other Pacific Islander; Non-Hispanic Pacific Islander |
| White/Caucasian | White; Caucasian; Caucasian/white; white/Caucasian; Non-Hispanic white; white Non-Hispanic; European white; European American; white British; Ethnically white British; Caucasian Ethnicity; European Ancestry; Nordic Origin |
| Two or More Races/Ethnicities | Mixed; Multi-Ethnic; Mixed Race; Multiracial; Bi-Racial; Mixed race (African American and Caucasian); Asian/Pacific Islander |
| Eurocentric | Non-white; Minority race (Black American or multi-race); Of Minority Race; other than Caucasian; Asian/Black/Mixed |
| Other | Other; Other/Multi-Race; Other race; Some other race |
| Not Reported/Unspecified | Did not specify; Unknown; Did not report |

Additionally, some papers reported other relevant sociodemographic descriptors, that were not race/ethnicity. These included papers that described the nationality or country of origin of participants within their sample, or information about the ethnic makeup of the community where the samples were recruited (see Supplementary Table 2).

To assess whether studies reported other common variables indicative of sociodemographic diversity, all studies were also binary-coded to indicate whether they did or did not report education and socioeconomic status/income. Reporting of education included years of education, highest qualification completed, and other bespoke groupings of educational bands (e.g., high-school, bachelors, graduate, and postgraduate degrees). Studies were coded as having reported a measure of socioeconomic status (SES)/income if they included information about income/household income bands (reported in various currencies), income-to-needs ratio, receipt of financial/public/housing assistance, employment status, housing status, or if the study used a quantitative measure of SES, or explicitly described their participants as low/middle/high SES, as low/middle/high income or low/middle/high socioeconomic background, as low/middle/upper class, or as living in low/middle/upper class neighborhoods.

## Results

This scoping review identified 185 total papers which met inclusion criteria. A summary table of these studies can be found listed in Supplementary Table 3. Included studies reported sample sizes ranging from 6 to 203 total participants (M=48.4, SD=32.8), for a total of 3,197 reported participants contributing to our understanding of the maternal brain. Studies were conducted across world regions (Sub-Saharan Africa, Middle East and North Africa, East Asia and Pacific, South Asia, Europe and Central Asia, and North America) and across 24 countries (Argentina, Australia, Belgium, Brazil, Cameroon, Canada, China, Czech Republic, Denmark, France, Germany, India, Israel, Italy, Japan, Kenya, Netherlands, South Africa, South Korea, Spain, Sweden, Switzerland, United Kingdom, and United States), with two multi-site studies conducted across regions (referred to here as ‘Intercontinental’). Papers were published between 2002 and 2023, and used structural (35 studies), task-based functional (98 studies), resting-state functional (36 studies), both task-based and resting-state functional (11 studies), and magnetic resonance spectroscopy (5 studies).

### Racial and ethnic representation in MRI studies of the maternal brain

Of the 185 papers, 71 papers (38.38%) reported race and/or ethnicity data, 107 papers (57.84%) studies reported no race or ethnicity data, and 7 papers (3.78%) reported a metric other than race and ethnicity (e.g., nationality/country of origin; Figure 2a). The 71 articles which reported racial and/or ethnic identities of participants included a total of 3,197 participants (Figure 2b). Of the total participants in studies that reported race and/or ethnicity data, participants were American Indian/American Native (*n*=7; 0.22%), Asian (*n*=390; 12.20%), Black/African American (*n*=391; 12.23%), Hispanic/Latine (*n*=246; 7.70%), Pacific Islander/Native Hawaiian (*n*=2; 0.06%), white/Caucasian (*n*=1497; 46.83%), and Two or More Races/ Ethnicities (*n*=21; 0.66%). An additional 13.23% of participants’ race and/or ethnicities were described in Eurocentric terms (e.g., “of minority race,” “non-white”; *n*=423), 1.72% of participants were described as “Other” (*n*=55), and 5.16% of participants’ race/ethnicities were “Not Reported” (*n*=165). 15.50% of the studies that reported race and/or ethnicity had a homogenous sample, in which all participants were from one racial/ethnic demographic. These homogenous studies took place in China, Germany, Sweden, the United Kingdom, and the United States. Of note, among the articles which report racial and/or ethnic identity, 50.70% (N=36 studies) include Hispanic/Latine identities in their demographics. Moreover, of the studies that report Hispanic/Latine identity, 83.3% (N=30 studies) report it in combination with race (Hispanic/Latine of any race), and 16.6% (N=6 studies) describe Hispanic/Latine identity separately as ethnicity, where 65 participants (23.90%) from these 6 studies were Hispanic/Latine, and 207 participants (76.10%) were non-Hispanic/Latine (Figure 3c). A subset of studies reporting race/ethnicity data used the information in their analyses as covariates or in direct group comparison (14 of 71 studies; 19.72%).

**Figure 2:**
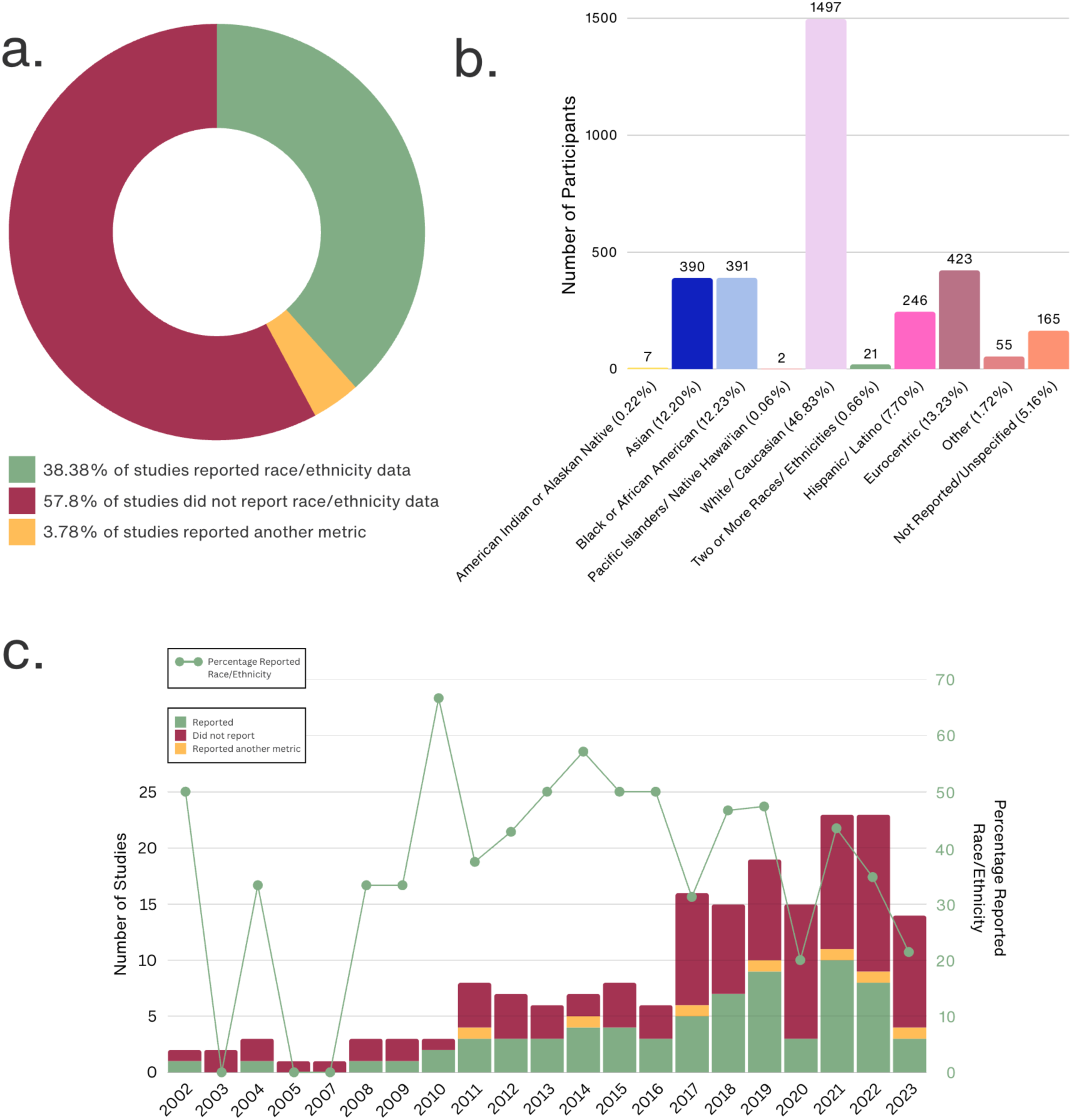
Reporting of Race and/or Ethnicity in Studies of the Maternal Brain. Fig2a. Percentage of studies which reported race and/or ethnicity demographics of their sample (shown in green), those that did not report any (shown in red), and those that reported another relevant metric (shown in yellow). Of those 38.38% of studies that reported race and/or ethnicity data, Fig2b. shows the number of participants for each reported racial/ethnic identity, sample size is shown on top of each bar, with percentage of total participants shown in axis labels. Fig 2c shows the reporting of race and/or ethnicity data by year. The bar chart shows the number of studies which reported race and/or ethnicity demographics of their sample (shown in green), those that did not report any race/ethnicity data (shown in red), and those that reported another relevant metric (shown in yellow) for each year. Percentage of papers reporting race/ethnicity data by year is overlayed as a line graph (shown in green).

**Figure 3:**
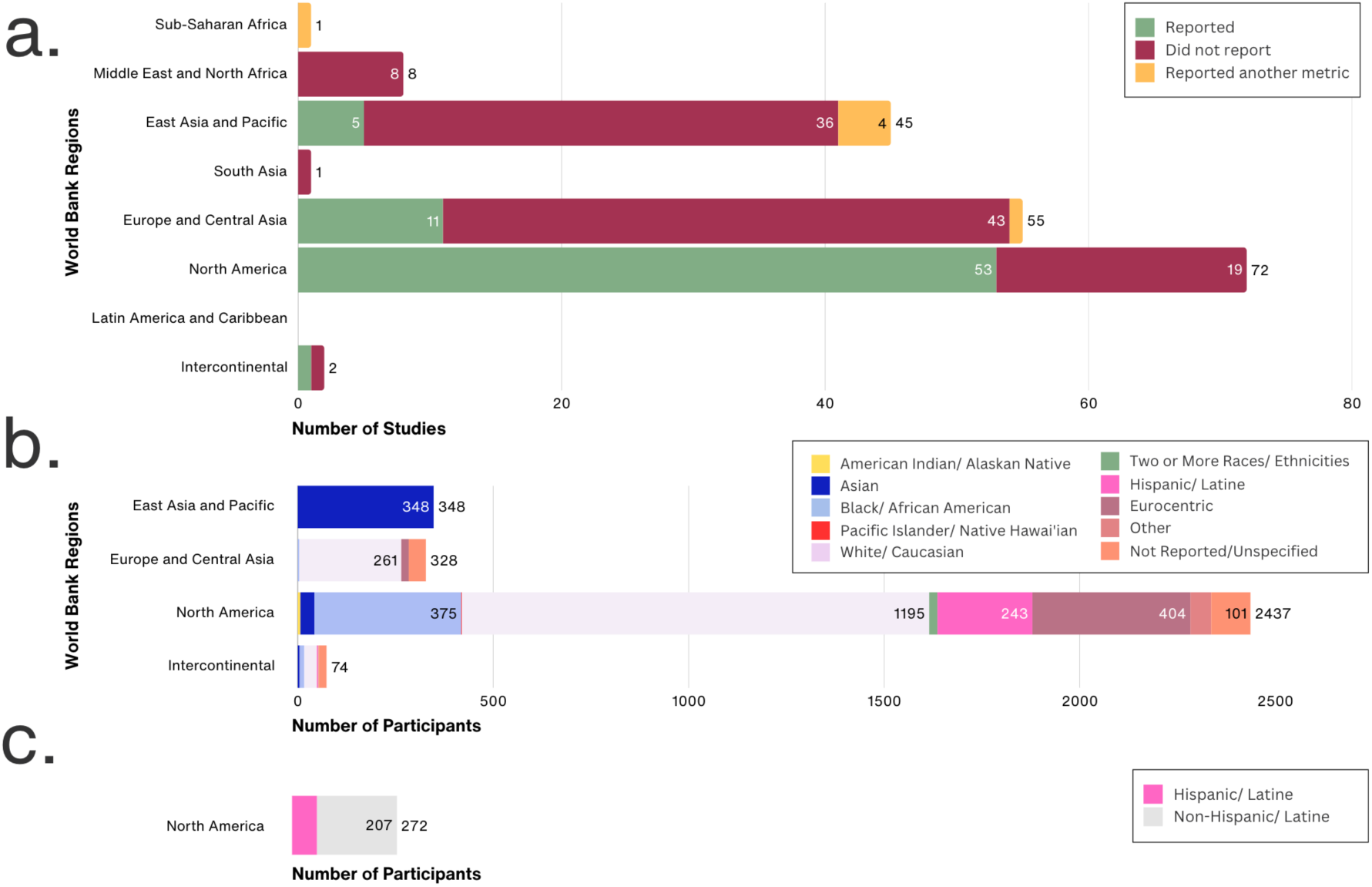
Racial and Ethnic Demographic Reporting by World Bank Region. Fig3a: Number of studies which reported race and/or ethnicity demographics of their sample (shown in green), those that did not report any race and/or ethnicity data (shown in red), and those that reported another relevant metric (shown in yellow) for each continent (Africa, Americas, Asia, Europe, Oceania, Intercontinental). The total number of studies from each continent is shown on top of each bar (in black), with the number reported/not reported/reported another metric within each stacked section. Of the studies that reported race and/or ethnicity data, Fig3b. shows the number of participants for each reported racial/ethnic identity for each continent. Fig 3c. shows the number of Hispanic/Latine (N=65) and non-Hispanic/Latine (N=207) identity in 6 studies from the United States which report Hispanic/Latine ethnicity in addition to race. N.B. some stacked bars are too thin to contain numbers, please refer to the results section where all sample sizes are listed in text.

The number of studies investigating the maternal brain using MRI has increased over time. In 2002, just two papers were published in this field. That number has increased 10-fold to a peak of 23 papers published in both 2021 and 2022. Despite the increase in publishing on the maternal brain using MRI, reporting of participant racial and ethnic data has remained largely consistent in a range of 20-57% across 17 of the last 21 years (Figure 2c; min 0% in 2003, 2005, and 2007, max 67% in 2010).

### Reporting of racial and ethnic diversity by continent

Reporting of racial and ethnic identity of participants varied by World Bank Region (Figure 3a). 0% of studies from Sub-Saharan Africa reported race and/or ethnicity (0/1 studies), 0% of studies from Middle East and North Africa (0/8 studies), 11.11% of studies from East Asia and Pacific regions (5/45 studies), 0% of studies from South Asia (0/1 studies), 20% of studies from Europe and Central Asia (11/55 studies), 73.61% of studies from North America (53/72 studies), and 50% of studies across multiple World Bank Regions (1/2 studies) reported any race and ethnicity data. 100% of studies from Sub-Sharan Africa (1/1 studies) reported another relevant metric (e.g. nationality/country of origin), 0% of studies from Middle East and North Africa (0/8 studies), 8.89% of studies from East Asia and Pacific regions (4/45 studies), 0% of studies from South Asia (0/1 studies), 1.82% of studies from Europe and Central Asia (1/55 studies), 0% of studies from North America (0/72 studies), and 0% of Intercontinental studies (0/2 studies). No studies were conducted in Latin America and the Caribbean. Of the total 3,197 participants with racial and/or ethnicity data reported across studies, 2,437 participants (76.23%) were from studies conducted in the United States.

Likewise, racial and ethnic diversity also varied across World Bank Regions (Figure 3b). In East Asia and Pacific regions, 100% of participants reported were Asian (348 of 348 participants). Of the 328 participants in studies from Europe and Central Asia, participants were Asian (*n*=1; 0.30%), Black/African-American (*n*=4; 1.18%), Eurocentric (*n*=19; 5.62%), white/Caucasian (*n*=270; 79.88%), and Not Reported (*n*=44; 13.02%). Of the 2,437 participants in studies from North America, participants were American Indian/Alaska Native (*n*=7; 0.29%), Asian (*n*=36; 1.48%), Black/African-American (*n*=375; 15.39%), Pacific Islander/Native Hawai’ian (*n*=2; 0.08%), Two or More Races/Ethnicities (*n*=21; 0.86%), Hispanic/Latine of any race (*n*=243; 9.97%), white/Caucasian (*n*=1195; 49.22%), Eurocentric (*n*=404; 16.58%), “Other” (*n*=53; 2.18%), and Not Reported (*n*=101; 4.14%). Of the 74 participants in studies conducted across multiple World Bank Regions, participants were Asian (*n*=5; 6.76%), Black/African-American (*n*=12; 16.22%), white/Caucasian (*n*=32; 43.24%), Hispanic/Latine of any race (*n*=3; 4.05%), “Other” (*n*=2; 2.70%), and Not Reported (*n*=20; 27.03%).

### Reporting of sociodemographic variables

To add perspective about reporting practices for other forms of sociodemographic diversity, we also identified the number of studies which reported SES/income and education. Of the 185 identified papers, 105 (56.76%) reported data describing education, and 70 (37.84%) reported data describing SES/income. This can be compared to 78 papers which reported race/ethnicity or another relevant metric (42.16%). 33 papers (17.84%) reported all three sociodemographic variables, and 45 papers (24.32%) reported none of the three. The overlaps between papers reporting race/ethnicity, education, and/or SES/income are shown in Figure 4.

**Figure 4.**
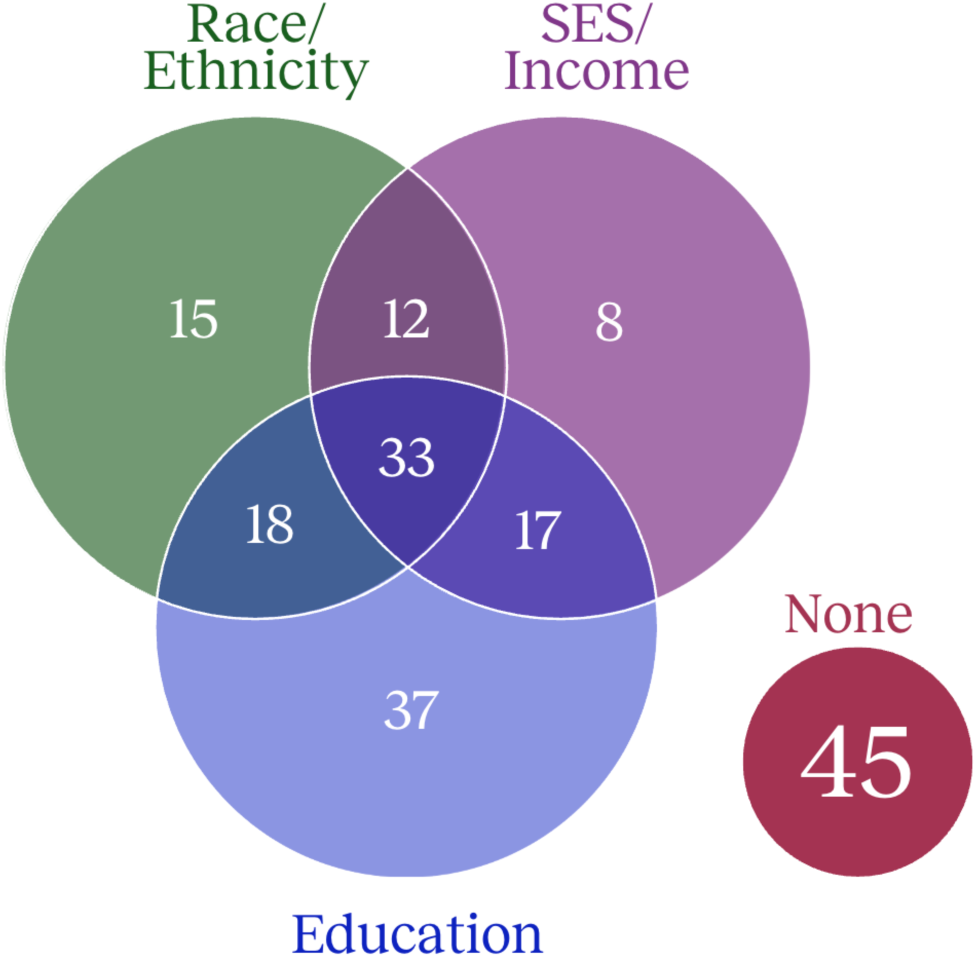
Reporting of Sociodemographic Variables. Number of studies which reported race/ethnicity demographics of their sample (shown in green), compared to the number which reported Education (shown in blue), and/or socioeconomic status (SES)/income (shown in purple), and those which reported none of these metrics (shown in dark pink).

## Discussion

Diverse and representative samples are essential for the integrity and generalizability of all scientific discoveries relevant to humans. However, current and historic systemic racism in neuroscience and the exclusion of historically minoritized identities limit our understanding of the human brain^10,12,13^, including research focused on motherhood^31^. As the field of parental neuroscience takes off, ensuring racial and ethnic diversity is a matter of both academic rigor and ethical imperative. Here, we add our voices to the growing demand for diversity, representation, and equity in neuroscience^10,12–14,16,17,41–48^, with a specific focus on racial and ethnic diversity in MRI studies of the maternal brain.

Our findings highlight three key issues in the study of the maternal brain using MRI: (1) the widespread underreporting of racial and ethnic demographics; (2) the overrepresentation of white participants; and (3) the overrepresentation of studies from North America and Europe and Central Asia. These findings raise concerns about the generalizability of existing research for the Global Majority – peoples who are Black, Indigenous, Asian, Latine, Middle Eastern, North African, Pacific Islander, and multiracial, who together comprise the majority of the world’s population. These findings also underscore the urgent need for concerted structural change in neuroscience research practices.

As we discuss these findings, we highlight the work and words of scholars who are leading diversity, equity, inclusion (DEI) and anti-racism efforts in neuroscience^10,12–14,16,17,41–48^, and point to existing resources which outline best practice guidelines for study design, recruitment, analyses, interpretation, and communication of results^13,48–50^.

### Underreporting of Racial and Ethnic Data in Maternal Brain Research

Of the 185 studies reviewed, just over one third (71 studies; 38.38%) report racial and/or ethnic demographics of their sample, representing a considerable underreporting of race and/or ethnicity in MRI studies of the maternal brain. This widespread omission in the maternal brain literature reflects a broader pattern in neuroscience, where racial and ethnic data are often overlooked or inconsistently reported^16^. Such underreporting limits the ability of researchers to assess and rectify biases and reinforces patterns of exclusion^10^. Further, we find that the underreporting of race and/or ethnicity data has been largely consistent across the last two decades, with the number of studies published on the maternal brain increasing over time, but the *percentage* of these studies reporting race and/or ethnicity remaining low (see Figure 1c). This suggests that reporting norms are not changing with the field, and concerted efforts are still required to change our practices moving forward.

Without transparent reporting of these data, assessing racial and/or ethnic representation across the literature is impossible. However, given that neuroscience research has historically relied on samples drawn from predominantly affluent white populations^13,15,16^, it is reasonable to assume that the 57.8% of studies that did not report these data likely overrepresent white participants, and/or contain largely homogenous samples, implicating the reproducibility and external validity of existing maternal brain findings for the Global Majority^15,51^.

Despite growing awareness, many scientific journals do not require researchers to disclose racial and/or ethnic demographics of their samples^10,16,52,53^. Without journal-mandated transparency, researchers face little obligation to assess sample diversity, allowing exclusionary practices to persist^10,17^. In attempts to combat this issue, the National Institutes of Health (NIH) in the United States now mandates that all NIH-funded clinical research studies report racial and/or ethnic demographics alongside sex and gender^54^. Some journals, such as *Nature*, have added race and/or ethnicity demographics into their reporting checklists and in turn, these reporting guidelines set a standard for scientific practice. The reporting of demographics remains critical to understand our sample populations and we strongly encourage the reporting of race and/or ethnicity data, even when recruiting homogenous samples (though see also ‘Global Disparities in Reporting Practices’ section for more nuance here). Transparent reporting practices support our understanding of representation for the field as a whole and provide the opportunity for accountability, self-reflection, and change.

### Overrepresentation of White Participants and Eurocentric Frameworks

Among studies that did report race and ethnicity data, nearly half of the participants were white/Caucasian, with Asian, Black/African American, Hispanic/Latine, Indigenous, Pacific Islander/Native Hawai’ian, and Middle East and North African participants underrepresented in MRI studies of the maternal brain. Our findings align with prior research demonstrating similar trends in other neuroimaging modalities. Penner and colleagues^31^ found that EEG studies of the maternal brain also disproportionately sampled white participants, where over half the participants were white/of European ancestry. Given this consistency between imaging modalities^31^, it is critical to acknowledge and address recruitment and reporting practices in maternal neuroscience before they become further entrenched in the field. The failure to recruit diverse participants has profound consequences for the validity of neuroscience research and suggests that our current understanding of the structure and function of the maternal brain may be biased and not generalizable to mothers of all racial and ethnic identities.

Moreover, the ways in which some racial and/or ethnic identities were reported in the studies reviewed here raise further concerns about inclusivity. In studies reporting race or ethnicity, one in five participants were categorized using descriptors that were either non-specific (e.g., ’Other,’ ‘not reported’) or reflective of Eurocentric frameworks (e.g., ‘non-white,’ ‘of minority race’). Describing participants in relation to whiteness reflects outdated and problematic frameworks which diminish the identities of historically marginalized groups by positioning whiteness as the default^13^. This lack of specificity renders almost a quarter of these data at best uninformative and, at worst, exclusionary, underscoring the need for more precise and inclusive racial and ethnic classifications in maternal brain research. One potential reason for the relatively high proportion of participants who self-selected/were designated ‘Other’ and/or ‘not reported’ may be the limited options available on the demographic forms used to record participants’ racial and ethnic identities. Whilst this is speculative, as we do not know how each study collected these data, many studies appeared to opt to align with the United States’ census guidelines, albeit with considerable variability (see Table 1). However, there are many identities which are not included, or are misrepresented, forcing people to either a.) choose between options that do not represent their identity, b.) designate themselves as ‘Other’, or c.) choose to ‘not describe’^13,55^. Capturing participants’ identities accurately and granularly provides richer and more meaningful data that can better describe literature on both an independent study and a global literature standpoint.

### Global Disparities in Reporting Practices

Our analysis also revealed geographic differences in race and/or ethnicity reporting across regions where studies were completed. While three quarters of studies that were conducted in North America report race and/or ethnicity, this was less common in studies from other regions, with 1 in 5 studies from Europe and Central Asia and 1 in 9 studies from East Asia and Pacific regions reporting race and/or ethnicity demographics. Furthermore, none of the studies from Sub-Saharan Africa, Middle East and North Africa, and South Asia reported race and/or ethnicity demographics.

Importantly, there are relevant regional, cultural, and legal differences in the collection and reporting of race and ethnicity data worldwide. In some European countries (e.g., Denmark, France, Germany, Sweden), collecting racial or ethnic data is legally prohibited and may appear insensitive or inappropriate to participants, rendering racial and ethnic transparency infeasible^13,49^. Further, it is important to acknowledge that the United States’ census reporting standards are not always relevant for other countries and the role of ‘race’ as it is understood in the American context does not adequately describe diversity globally. In some countries/regions, especially where race is more homogenous, ethnicity and other sociodemographic variables may become more salient or more informative for describing one’s sample, including information regarding socioeconomic status, migration status, language spoken at home, religious or cultural affiliation, geographical location, or any other factor which may contribute to the experience of social inequity, discrimination, or persecution^49^. When this is the case, we encourage researchers to report these types of demographics for better understanding of their populations sample. Generally, in neuroscience, socioeconomic status and education are also collected and reported. However, within the MRI maternal brain literature, these too were underreported and require reflection and consideration for reporting standards within the maternal brain field moving forward – with 24.32% of studies reporting neither metric of race and/or ethnicity nor socioeconomic status and/or education. Furthermore, we found that 3.78% of studies reported another metric intending to convey identity, without specifically reporting race and/or ethnicity of their sample. For example, nationality, country of origin, and ethnic makeup of the community where the sample were recruited were used in place of race and/or ethnicity in some studies from Europe and Central Asia, East Asia and Pacific regions, and Sub-Saharan Africa, underscoring that the most important variables to report are often highly specific to population and context.

Additionally, of the studies that reported race and/or ethnicity, 15.50% included a homogenous sample, in which all participants within those studies shared the same racial and/or ethnic identity. Whilst the recruitment of a homogenous sample is sometimes difficult to avoid, for example, when living in a highly homogenous country and/or region, we encourage researchers to describe the sociodemographic variables most relevant to understanding diversity within the context of their population^15,42,48^.

When stratifying by geographic region, the disproportionate distribution of studies becomes apparent. Of the included studies, 72 were conducted in North America and 55 in Europe and Central Asia, whilst only 11 studies were conducted in Sub-Saharan Africa, Middle East and North Africa, and South Asia combined, and no studies conducted in Latin America and the Caribbean. One stark global disparity that contributes to these findings is the profoundly unequal access to research-grade MRI scanners across the globe. It is estimated that 66% of the world does not have access to MRI scanners, disproportionately impacting lower and middle-income countries, and regional and rural areas, with significant barriers including high costs of equipment acquisition and maintenance, shortages of skilled technicians and researchers, and scarcer research funding and infrastructure^56–58^. Whilst there are recent efforts to make MRI portable and more globally accessible^59,60^, these disparities contribute to a lack of global representation in neuroimaging research, limiting the generalizability of findings and reinforcing existing inequities related to race and/or ethnicity.

### De-centering white/Western Bias from the Study of Motherhood

Whilst the experience of pregnancy, birth, and lactation are biological processes for gestational parents, it is important to consider that motherhood is fundamentally a social relationship and is heavily influenced by culture and context. Researchers should therefore reflect on whether and how they are applying an ethnocentric lens to the study of the maternal brain, and ensure that white motherhood, and Western ideals of parenting behavior are not inadvertently centered as ‘normal,’ or ‘universal’^11,61,62^. In addition to reporting the demographics of samples tested, researchers should consider how culture and context constrain interpretations, particularly when examining ‘maternal behavior,’ as the behaviors constituting adaptive parenting vary across different groups based on both cultural and sociocontextual factors^61,63^. Although made up of a variety of cultures and ethnicities, Western and predominantly white interpretations of maternal behavior are often used to design research studies, and behavioral and psychometric tests. Researchers should critically evaluate whether their study designs, hypotheses, and interpretations reflect a Western-centric perspective and, where applicable, integrate culturally diverse frameworks. Without this reflexivity, the field risks reinforcing narrow assumptions about motherhood, maternal behavior, and brain function that do not translate across different sociocultural contexts. Expanding theoretical models and methodological approaches will enhance the relevance and applicability of maternal brain research on a global scale.

It should go without saying that diverse global samples can additionally provide essential mechanistic insight into the neural changes related to motherhood. For example, diverse samples allow the investigation of whether maternal brain changes may be universal and/or culturally or regionally specific, and whether they relate to social factors including gendered norms of caregiving, parental leave policy, intergenerational support, and alloparental care. Uncovering answers to these questions can guide policy change and inform equitable clinical care for myriad caregiver contexts and populations, but require careful, considerate investigation of evidence collected from diverse global samples.

### A Call for Structural Change

Our study focuses on the reporting of race and ethnicity data, yet true inclusivity and equity in neuroscience research requires deeper engagement beyond demographic documentation. The barriers to recruiting representative samples, along with strategies to combat exclusionary practices, have been extensively outlined in prior work^10,12–14,16,17,41–48^. In addition to generalized recommendations^49^, the unique challenges of studying mothers^31^, issues specific to MRI methodology^42^, and the importance of considering intersectional identity^48^ have been elegantly detailed, and we urge researchers to read, reflect on, and implement the recommendations within these important works. As Cardenas-Iniguez et al. (2024) emphasize, considerations of race and ethnicity must be integrated at every stage of research—from study conceptualization to data analysis, interpretation, and communication of results^13^.

In maternal brain research, it is particularly critical to recognize the structural and socio-contextual factors influencing participation, especially for mothers from historically marginalized communities. Mothers face significant logistical and financial burdens when engaging in research, including the need to arrange childcare, which can be costly and inconvenient. These challenges disproportionately impact low-income participants, single mothers, and those reliant on public transportation, further exacerbating disparities in study participation^23,24^. Additionally, many maternal brain studies require mothers to be separated from their child for extended periods for cognitive or behavioral tasks and/or neuroimaging, which can create emotional distress for both mother and child, beyond the financial and logistical costs of participation. Without addressing these systemic barriers, the field will continue to struggle with under-recruitment of racially and ethnically diverse samples and perpetuate deepening health inequities.

The intersection of pregnancy, early motherhood, and healthcare research is particularly fraught given the historical and ongoing medical discrimination against Black women and other marginalized groups. While reluctance to participate in research is often attributed to past injustices^64^, recent work^65^ highlights that contemporary healthcare experiences continue to profoundly shape health-related decision-making. Rather than accepting distrust as a static barrier, it is the responsibility of researchers to actively mend relationships with communities, foster positive and reciprocal relationships, and engage in transparent, culturally sensitive communication with participants and communities. The current trajectory of maternal brain research risks perpetuating exclusionary practices that compromise the field’s scientific rigor and ethical standing. As a research community, we have a shared responsibility and accountability for ensuring that future research is built on diverse and representative samples.

### Limitations

The maternal brain literature is a small but growing field, and many research groups have published more than one paper from the same sample/data collection effort, reflecting the difficulties and costs involved in collecting these precious data. This means that some of the same participants are included in the analyses of multiple papers, with the number of unique participants lower than those reported here. As it is unclear in many cases which participants were and were not included across multiple studies, and subsequently difficult to accordingly adjust race and/or ethnicity data, we chose to treat each paper as independent for our coding purposes. Our understanding of the maternal brain is somewhat further limited to these repeated participants, and therefore the diversity and representation (or lack thereof) in these studies is made more influential. Additionally, the number of participants and their racial and/or ethnic identities reported here also include the participants comprising control/comparison groups, as the racial and/or ethnic identities of these participants also contribute to our understanding of the maternal brain. Additionally, as our inclusion criteria included English language and published works, it is possible that relevant papers published in languages other than English, or that were unpublished may have increased the diversity and representation beyond what was reported here. Finally, we largely defined categories based on the United States’ census guidelines. As noted by Cardenas-Iniguez et al., “*Globally, harmonizing these categories can be complex, as countries differ in how they measure the diversity of their population… Researchers should particularly refrain from assuming that race and ethnicity categories have identical meanings globally”* (*p*. 618). Whilst the majority of participants with reported race and/or ethnicity data were from studies in the United States, and the categories appeared sufficient to represent the reported data, it is possible that by aggregating the data by the United States’ census categories we have misrepresented some participants’ identities, though this was done with great care, discussion, and transparency (see Table 2).

## Conclusion

Our findings highlight three key issues in the study of the maternal brain using MRI: (1) the widespread underreporting of racial and ethnic demographics; (2) the overrepresentation of white participants; and (3) the limited distribution of study locations across the globe. These findings reflect a persistent WEIRD (Western, Educated, Industrialized, Rich, and Democratic)^15^ bias in neuroscience, where the majority of research participants are drawn from homogenous, high-income populations that do not reflect global diversity. This bias has critical implications for the generalizability of the data and our overall understanding of the maternal brain.

Women’s health is receiving a recent surge in attention and neuroscience research focusing on factors disproportionately impacting women is booming. However, as the wave of women’s health rises, scientists must highlight and champion intersectionality, diversity, and representation, and be ever cautious of participating in and perpetuating “white feminism.” In order for our research to improve the lives of women and birthing people, it must include and represent more people and communities. Without significant changes in reporting standards, recruitment practices, and methodological approaches, the field risks reinforcing the same systemic biases that have historically marginalized minoritized populations in neuroscience research. Because these efforts are vital, yet often costly and inconvenient, they must be actively prioritized in budgeting, recruitment, and study design, and not treated as an afterthought.

Lack of representative samples contribute to a cycle of failed replicability and generalizability in neuroscience^16^. An essential step to understanding this field and who it impacts, is through the accurate and appropriate reporting of sample demographics. Without this knowledge, there is a risk of reenacting the replication crisis^66,67^ in psychology, positioning researchers at risk of having to redo this research again in 20 years. This scoping review echoes the existing chorus of voices that have championed diversity and representation in neuroscience^10,12–14,16,17,41–48,50^ and showcase the importance of these efforts for studies of the maternal brain.

## Acknowledgements

E.R.O is supported by a Yale Kavli Institute for Neuroscience Postdoctoral Fellowship and an American Association for University Women International Postdoctoral Fellowship. C.M is supported by a Yale Kavli Institute for Neuroscience Postdoctoral Award for Academic Diversity and Yale Science Fellowship. K.W and K.A are supported by T32 NS041228, K.W is supported by F31 DA059248. This publication was made possible by CTSA Grant Number TL1 TR001864 (K.A support) from the National Center for Advancing Translational Science (NCATS), a component of the National Institutes of Health (NIH). Its contents are solely the responsibility of the authors and do not necessarily represent the official views of NIH. J.A.R. is supported by the Stanford University Knight–Hennessy Scholars Program, as well as the National Academies of Sciences, Engineering and Medicine’s Ford Foundation Predoctoral Fellowship. F.P was supported by F32 DA055389. T.Y was supported by a Yale Kavli Institute for Neuroscience Postdoctoral Fellowship. H.J.V.R is supported by R01 HD108218 and R01 DA050636. The authors would like to thank Vasean Daniels and Vermetha Polite of the Cushing/Whitney Medical Library for technical support.

