## Supplementary Materials for "Scanning for Representation: A Scoping Review of Racial and Ethnic Diversity in MRI Studies of the Maternal Brain"

### Supplementary Table 1: Search Strategy

Ovid MEDLINE(R) ALL <1946 to November 03, 2022>

| | |
|---|---|
| 1 | postpartum period/ or (postpartum or post partum or puerperium or puerperal or pregnan* or mother*).tw,kw. or ((after or following or post or giving or give or gave) adj2 (birth or childbirth)).tw,kw. or ((mother* or maternal) adj3 (postnatal or peripartum or perinatal)).tw,kw. |
| 2 | exp functional neuroimaging/ or (neuroimaging or functional connect*).tw,kw. or ((BOLD or neural or brain or grey matter or gray matter or white matter or cortical thickness) adj3 (plasti* or activ* or increas* or decreas* or chang* or reduc* or affect* or adapt*)).tw,kw. |
| 3 | 1 and 2 |
| 4 | 3 not (animals/ not (animals/ and humans/)) |

### Supplementary Table 2: Descriptions of seven papers reporting another relevant metric

| **Metric reported** | **Verbatim Description** | **Country** | **World Bank Region** |
|---|---|---|---|
| Demographics of the community where the sample was collected | "Pregnant women were recruited from two primary health care clinics, Mbekweni (serving a predominantly black African community) and TC Newman (Serving a mixed ancestry community)." | South Africa | Sub-Saharan Africa |
| Nationality | "Japanese parents" | Japan | East Asia and Pacific |
| Nationality | "Japanese parents" | Japan | East Asia and Pacific |
| Nationality | "All participants were native Koreans" | South Korea | East Asia and Pacific |
| Nationality/ Cultural Background | The fMRI data was acquired on mothers from "three distinct cultures": The United States sample included "European American mothers". The China sample included "Shanghai Chinese mothers". The Italy sample included "Mothers and Non-Mothers in Italy", referred to as "Italian Mothers". | Argentina, Belgium, Brazil, Cameroon, France, Kenya, Israel, Italy, Japan, South Korea, and United States (N.B. *fMRI results only from samples in United States, China, and Italy*). | Intercontinental |
| Nationality/ Cultural Background | Refer to their sample in the Introduction and Discussion, as "Chinese mothers/women" (discuss their results compared to "Western mothers" of previous studies, for instance). | China | East Asia and Pacific |
| Country of Origin | "In the postpartum group, 63 (80.8%) were Germans, 2 (2.6%) were from other West European countries (Belgium, Netherlands), 1 (1.3%) was from Spain, 8 (10.3%) were from Eastern Europe (Poland, Lithuania, Russia, Slovakia), and 4 (5.1%) were from African, Arab, or Asian countries (Morocco, Iran, Liberia, Tajikistan). In the nulliparous control group, 26 participants (70.3%) were from Germany, 3 (8.1%) from a West European country (France), 3 (8.1%) from the Mediterranean region (Spain, Greece, Turkey), 4 (10.8%) from Eastern Europe (Russia, Bulgaria), and 1 (2.7%) from Africa (no specific country was mentioned). Three nulliparous participants did not provide information regarding their country of origin." | Germany | Europe and Central Asia |

### Supplementary Table 3: Included Papers

| Author | Year | Title | DOI | Country | World Bank Region |
|---|---|---|---|---|---|
| Abel | 2018 | Preliminary evidence for neural responsiveness to infants in mothers with schizophrenia and the implications for healthy parenting | https://doi.org/10.1016/j.schres.2017.11.033 | United Kingdom | Europe and Central Asia |
| Abraham | 2014 | Father's brain is sensitive to childcare experiences | www.pnas.org/cgi/doi/10.1073/pnas.1402569111 | Israel | Middle East and North Africa |
| Abraham | 2017 | The Human Coparental Bond Implicates Distinct Corticostriatal Pathways: Longitudinal Impact on Family Formation and Child Well-Being | doi: 10.1038/npp.2017.71 | Israel | Middle East and North Africa |
| Abraham | 2018 | Empathy networks in the parental brain and their long-term effects on children's stress reactivity and behavior adaptation | http://dx.doi.org/10.1016/j.neuropsychologia.2017.04.015 | Israel | Middle East and North Africa |
| An | 2021 | Decreased grey matter volumes in unaffected mothers of individuals with autism spectrum disorder reflect the broader autism endophenotype | https://doi.org/10.1038/s41598-021-89393-z | Japan | East Asia Pacific |
| Aran | 2023 | Neural activation to infant cry among Latina and non-Latina White mothers | https://doi.org/10.1016/j.bbr.2023.114298 | USA | North America |
| Atzil | 2011 | Specifying the neurobiological basis of human attachment: brain, hormones, and behavior in synchronous and intrusive mothers | 10.1038/npp.2011.172 | Israel | Middle East and North Africa |
| Atzil | 2012 | Synchrony and specificity in the maternal and the paternal brain: relations to oxytocin and vasopressin | http://dx.doi.org/10.1016/j.jaac.2012.06.008 | Israel | Middle East and North Africa |
| Atzil | 2014 | The brain basis of social synchrony | https://doi.org/10.1093/scan/nst105 | Israel | Middle East and North Africa |
| Atzil | 2017 | Dopamine in the medial amygdala network mediates human bonding | www.pnas.org/cgi/doi/10.1073/pnas.1612233114 | USA | North America |
| Bak | 2021 | Neural correlates of empathy for babies in postpartum women: A longitudinal study | 10.1002/hbm.25435 | South Korea | East Asia Pacific |
| Bannbers | 2013 | Prefrontal activity during response inhibition decreases over time in the postpartum period | http://dx.doi.org/10.1016/j.bbr.2012.12.003 | Sweden | Europe and Central Asia |
| Barrett | 2012 | Maternal affect and quality of parenting experiences are related to amygdala response to infant faces | ttps://doi.org/10.1080/17470919.2011.609907 | Canada | North America |
| Bartels | 2004 | The neural correlates of maternal and romantic love | doi:10.1016/j.neuroimage.2003.11.003 | United Kingdom | Europe and Central Asia |
| Bjertrup | 2021 | Neurocognitive processing of infant stimuli in mothers and non-mothers: psychophysiological, cognitive and neuroimaging evidence | doi: 10.1093/scan/nsab002 | Denmark | Europe and Central Asia |
| Bjertrup | 2022 | Reduced prefrontal cortex response to own vs. unknown emotional infant faces in mothers with bipolar disorder | https://doi.org/10.1016/j.euroneuro.2021.09.011 | Denmark | Europe and Central Asia |
| Bojja | 2018 | Clinical and imaging profile of patients with new-onset seizures & a presumptive diagnosis of eclampsia - A prospective observational study | https://doi.org/10.1016/j.preghy.2018.02.008 | India | South Asia |
| Bornstein | 2017 | Neurobiology of culturally common maternal responses to infant cry | www.pnas.org/cgi/doi/10.1073/pnas.1712022114 | Argentina, Belgium, Brazil, Cameroon, France, Kenya, Israel, Italy, Japan, South Korea, and USA | Intercontinental |
| Bublitz | 2022 | Maternal History of Childhood Maltreatment and Brain Responses to Infant Cues Across the Postpartum Period | 10.1177/10775595221128952 | USA | North America |

| | | | | | |
|---|---|---|---|---|---|
| Capistrano | 2022 | Maternal socioeconomic disadvantage, neural function during volitional emotion regulation, and parenting | https://doi.org/10.1080/17470919.2022.2082521 | USA | North America |
| Carmona | 2019 | Pregnancy and adolescence entail similar neuroanatomical adaptations: A comparative analysis of cerebral morphometric changes | 10.1002/hbm.24513 | Spain | Europe and Central Asia |
| Chao | 2020 | Severe pre-eclamptic women with headache: is posterior reversible encephalopathy syndrome an associated concurrent finding?+C26:AO26 | https://doi.org/10.1186/s12884-020-03017-4 | China | East Asia Pacific |
| Chase | 2014 | Disrupted posterior cingulate-amygdala connectivity in postpartum depressed women as measured with resting BOLD fMRI | 10.1093/scan/nst083 | USA | North America |
| Che | 2020 | Altered Spontaneous Neural Activity in Peripartum Depression: A Resting-State Functional Magnetic Resonance Imaging Study | 10.3389/fpsyg.2020.00656 | China | East Asia Pacific |
| Chechko | 2022 | The expectant brain-pregnancy leads to changes in brain morphology in the early postpartum period | https://doi.org/10.1093/cercor/bhab463 | Germany | Europe and Central Asia |
| Chechko | 2023 | Neural responses to monetary incentives in postpartum women affected by baby blues | https://doi.org/10.1016/j.psyneuen.2022.105991 | Germany | Europe and Central Asia |
| Chen | 2023 | Aberrant structural and functional alterations in postpartum depression: a combined voxel-based morphometry and resting-state functional connectivity study | 10.3389/fnins.2023.1138561 | China | East Asia Pacific |
| Cheng | 2020 | Regional cerebral activity abnormality in pregnant women with antenatal depression | https://doi.org/10.1016/j.jad.2020.05.107 | China | East Asia Pacific |
| Cheng | 2021 | Cerebral regional homogeneity alternation of pregnant women with antenatal depression during the pandemic | doi: 10.3389/fpsyt.2021.627871 | China | East Asia Pacific |
| Cheng | 2022 | Abnormal dynamics of resting-state functional activity and couplings in postpartum depression with and without anxiety | https://doi.org/10.1093/cercor/bhac038 | China | East Asia Pacific |
| Cheng | 2022 | Altered functional connectivity density and couplings in postpartum depression with and without anxiety | https://doi.org/10.1093/scan/nsab127 | China | East Asia Pacific |
| Cheng | 2022 | Prolactin mediates the relationship between regional gray matter volume and postpartum depression symptoms | https://doi.org/10.1016/j.jad.2022.01.051 | China | East Asia Pacific |
| Cheng | 2022 | Social support mediates the influence of cerebellum functional connectivity strength on postpartum depression and postpartum depression with anxiety | https://doi.org/10.1038/s41398-022-01781-9 | China | East Asia Pacific |
| Chu | 2022 | Pregnancy leads to changes in the brain functional network: a connectome analysis | 0.1007/s11682-021-00561-1 | China | East Asia Pacific |
| Deligiannidis | 2013 | GABAergic neuroactive steroids and resting-state functional connectivity in postpartum depression: a preliminary study | 10.1016/j.jpsychires.2013.02.010. | USA | North America |
| Deligiannidis | 2019 | Resting-state functional connectivity, cortical GABA, and neuroactive steroids in peripartum and peripartum depressed women: a functional magnetic resonance imaging and spectroscopy study | https://doi.org/10.1038/s41386-018-0242-2 | USA | North America |
| Dong | 2022 | Altered dynamic amplitude of low-frequency fluctuations in patients with postpartum depression | https://doi.org/10.1016/j.bbr.2022.113980 | China | East Asia Pacific |
| Dudin | 2019 | Amygdala and affective responses to infant pictures: Comparing depressed and non-depressed mothers and non-mothers | 10.1111/jne.12790 | Canada | North America |
| Dufford | 2019 | Maternal brain resting-state connectivity in the postpartum period | doi:10.1111/jne.12737 | USA | North America |
| Eckstein | 2019 | The NeMo real-time fMRI neurofeedback study: protocol of a randomised controlled clinical intervention trial in the neural foundations of mother-infant bonding | 10.1136/bmjopen-2018-027747 | Germany | Europe and Central Asia |

| | | | | | |
|---|---|---|---|---|---|
| Elmadih | 2016 | Natural variation in maternal sensitivity is reflected in maternal brain responses to infant stimuli | http://dx.doi.org/10.1037/bne0000161 | United Kingdom | Europe and Central Asia |
| Finnegan | 2021 | Mothers' neural response to valenced infant interactions predicts postpartum depression and anxiety | https://doi.org/10.1371/journal.pone.0250487 | USA | North America |
| Firk | 2018 | Down-regulation of amygdala response to infant crying: A role for distraction in maternal emotion regulation | http://dx.doi.org/10.1037/emo0000373 | Germany | Europe and Central Asia |
| Ghuman | 2022 | Prospective Investigation of Glutamate Levels and Percentage Gray Matter in the Medial Prefrontal Cortex in Females at Risk for Postpartum Depression | 10.2174/1570159X20666220302101115 | Canada | North America |
| Gingnell | 2015 | Emotion Reactivity Is Increased 4-6 Weeks Postpartum in Healthy Women: A Longitudinal fMRI Study | 10.1371/journal.pone.0128964 | Sweden | Europe and Central Asia |
| Gingnell | 2017 | Emotional anticipation after delivery - a longitudinal neuroimaging study of the postpartum period | 10.1038/s41598-017-00146-3 | Sweden | Europe and Central Asia |
| Giuliani | 2019 | A Preliminary Study Investigating Maternal Neurocognitive Mechanisms Underlying a Child-Supportive Parenting Intervention | 10.3389/fnbeh.2019.00016 | USA | North America |
| Grande | 2021 | Postpartum Stress and Neural Regulation of Emotion among First-Time Mothers | https://doi.org/10.3758/s13415-021-00914-9 | USA | North America |
| Gregory | 2015 | Oxytocin increases VTA activation to infant and sexual stimuli in nulliparous and postpartum women | http://dx.doi.org/10.1016/j.yhbeh.2014.12.009 | USA | North America |
| Guo | 2018 | Severity of anxiety moderates the association between neural circuits and maternal behaviors in the postpartum period | 10.3758/s13415-017-0516-x | USA | North America |
| Hipwell | 2015 | Right Frontoinsular Cortex and Subcortical Activity to Infant Cry Is Associated with Maternal Mental State Talk | DOI:10.1523/JNEUROSCI.1286-15.2015 | USA | North America |
| Ho | 2017 | Depression alters maternal extended amygdala response and functional connectivity during distress signals in attachment relationship | 10.1016/j.bbr.2017.02.045 | USA | North America |
| Ho | 2020 | Potential Neural Mediators of Mom Power Parenting Intervention Effects on Maternal Intersubjectivity and Stress Resilience | 10.3389/fpsyt.2020.568824 | USA | North America |
| Hoekzema | 2017 | Pregnancy leads to long-lasting changes in human brain structure | 10.1038/nn.4458 | Spain | Europe and Central Asia |
| Hoekzema | 2020 | Becoming a mother entails anatomical changes in the ventral striatum of the human brain that facilitate its responsiveness to offspring cues | https://doi.org/10.1016/j.psyneuen.2019.104507 | Spain | Europe and Central Asia |
| Hoekzema | 2022 | Mapping the effects of pregnancy on resting state brain activity, white matter microstructure, neural metabolite concentrations and grey matter architecture | https://doi.org/10.1038/s41467-022-33884-8 | Netherlands | Europe and Central Asia |
| Holdcroft | 2005 | Phosphorus-31 brain MR spectroscopy in women during and after pregnancy compared with nonpregnant control subjects | DOI missing, pubmed link: https://pubmed.ncbi.nlm.nih.gov/15709134/ | United Kingdom | Europe and Central Asia |
| Houtchens | 2020 | MRI activity in MS and completed pregnancy: Data from a tertiary academic center | 10.1212/NXI.0000000000000890 | USA | North America |
| Huang | 2023 | Structural and functional improvement of amygdala sub-regions in postpartum depression after acupuncture | doi: 10.3389/fnhum.2023.1163746 | China | East Asia Pacific |
| Kersting | 2009 | Neural activation underlying acute grief in women after the loss of an unborn child | 10.1176/appi.ajp.2009.08121875 | Germany | Europe and Central Asia |

| | | | | | |
|---|---|---|---|---|---|
| Kim | 2010 | Perceived quality of maternal care in childhood and structure and function of mothers' brain | 10.1111/j.1467-7687.2009.00923.x | USA | North America |
| Kim | 2010 | The plasticity of human maternal brain: longitudinal changes in brain anatomy during the early postpartum period | 10.1037/a0020884 | USA | North America |
| Kim | 2011 | Breastfeeding, brain activation to own infant cry, and maternal sensitivity | 10.1111/j.1469-7610.2011.02406.x | USA | North America |
| Kim | 2014 | Mothers' unresolved trauma blunts amygdala response to infant distress | https://doi.org/10.1080/17470919.2014.896287 | USA | North America |
| Kim | 2015 | A Prospective Longitudinal Study of Perceived Infant Outcomes at 18-24 Months: Neural and Psychological Correlates of Parental Thoughts and Actions Assessed during the First Month Postpartum | doi: 10.3389/fpsyg.2015.01772 | USA | North America |
| Kim | 2016 | Socioeconomic disadvantages and neural sensitivity to infant cry: role of maternal distress | 10.1093/scan/nsw063 | USA | North America |
| Kim | 2017 | Mothers with substance addictions show reduced reward responses when viewing their own infant's face | 10.1002/hbm.23731 | USA | North America |
| Kim | 2017 | Socioeconomic disadvantage, neural responses to infant emotions, and emotional availability among first-time new mothers | doi:10.1016/j.bbr.2017.02.001 | USA | North America |
| Kim | 2018 | Cortical thickness variation of the maternal brain in the first 6 months postpartum: associations with parental self-efficacy | https://doi.org/10.1007/s00429-018-1688-z | USA | North America |
| Kim | 2020 | Associations between stress exposure and new mothers' brain responses to infant cry sounds | doi:10.1016/j.neuroimage.2020.117360 | USA | North America |
| Kim | 2022 | Intergenerational neuroimaging study: mother-infant functional connectivity similarity and the role of infant and maternal factors | https://doi.org/10.1093/cercor/bhab408 | USA | North America |
| Kim | 2022 | Trait coping styles and the maternal neural and behavioral sensitivity to an infant | doi.org/10.1038/s41598-022-18339-w | USA | North America |
| Koopowitz | 2023 | PTSD and comorbid MDD is associated with activation of the right frontoparietal network | https://doi.org/10.1016/j.pscychresns.2023.111630 | South Africa | Sub-Saharan Africa |
| Kowalczyk | 2021 | Neurocognitive correlates of working memory and emotional processing in postpartum psychosis: an fMRI study | https://doi.org/10.1017/S0033291720000471 | United Kingdom | Europe and Central Asia |
| Kurosaki | 2018 | Effects of perinatal blood pressure on maternal brain functional connectivity | https://doi.org/10.1371/journal.pone.0203067 | Japan | East Asia Pacific |
| Landi | 2011 | Maternal neural responses to infant cries and faces: relationships with substance use | 10.3389/fpsyt.2011.00032 | USA | North America |
| Laurent | 2011 | Neural correlates of hypothalamic-pituitary-adrenal regulation of mothers with their infants | doi:10.1016/j.biopsych.2011.06.011 | USA | North America |
| Laurent | 2012 | A cry in the dark: depressed mothers show reduced neural activation to their own infant's cry | doi:10.1093/scan/nsq091 | USA | North America |
| Laurent | 2012 | The missing link: mothers' neural response to infant cry related to infant attachment behaviors | http://dx.doi.org/10.1016/j.infbeh.2012.07.007 | USA | North America |
| Laurent | 2013 | A face a mother could love: depression-related maternal neural responses to infant emotion faces | doi:10.1080/17470919.2012.762039 | USA | North America |
| Lehmann | 2021 | Brain MRI activity during the year before pregnancy can predict post-partum clinical relapses | 10.1177/13524585211002719 | Israel | Middle East and North Africa |
| Lenzi | 2009 | Neural basis of maternal communication and emotional expression processing during infant preverbal stage | 10.1093/cercor/bhn153 | Italy | Europe and Central Asia |

| | | | | | |
|---|---|---|---|---|---|
| Lenzi | 2016 | Mothers with depressive symptoms display differential brain activations when empathizing with infant faces | http://dx.doi.org/10.1016/j.pscychresns.2016.01.019 | Italy | Europe and Central Asia |
| Leon | 2019 | Limbic-visual attenuation to crying faces underlies neglectful mothering | https://doi.org/10.1038/s41598-019-42908-1 | Spain | Europe and Central Asia |
| Leon | 2021 | Distinctive Frontal and Occipitotemporal Surface Features in Neglectful Parenting | https://doi.org/10.3390/brainsci11030387 | Spain | Europe and Central Asia |
| Li | 2019 | Altered Functional Connectivity and Brain Network Property in Pregnant Women With Cleft Fetuses | doi: 10.3389/fpsyg.2019.02235 | China | East Asia Pacific |
| Li | 2021 | Abnormalities of cortical structures in patients with postpartum depression: A surface-based morphometry study | https://doi.org/10.1016/j.bbr.2021.113340 | China | East Asia Pacific |
| Li | 2023 | Aberrant resting-state regional activity in patients with postpartum depression | 10.3389/fnhum.2022.925543 | China | East Asia Pacific |
| Lisofsky | 2016 | Differences in navigation performance and postpartal striatal volume associated with pregnancy in humans | http://dx.doi.org/10.1016/j.nlm.2016.08.0221074-7427/ | Germany | Europe and Central Asia |
| Lisofsky | 2019 | Postpartal Neural Plasticity of the Maternal Brain: Early Renormalization of Pregnancy-Related Decreases? | DOI: 10.33594/000000105 | Germany | Europe and Central Asia |
| Long | 2023 | Altered MRI Diffusion Properties of the White Matter Tracts Connecting Frontal and Thalamic Brain Regions in First-Episode, Drug-Naive Patients With Postpartum Depression | 10.1002/jmri.28346 | China | East Asia Pacific |
| Lorberbaum | 2002 | A potential role for thalamocingulate circuitry in human maternal behavior | https://doi.org/10.1016/S0006-3223(01)01284-7 | USA | North America |
| Lord | 2012 | Stress response in postpartum women with and without obsessive-compulsive symptoms: an fMRI study | 10.1503/jpn.110005 | Canada | North America |
| Lorenz | 2019 | Neural correlates of emotion processing comparing antidepressants and exogenous oxytocin in postpartum depressed women: An exploratory study | https://doi.org/10.1371/journal.pone.0217764 | USA | North America |
| Luders | 2018 | Potential Brain Age Reversal after Pregnancy: Younger Brains at 4-6 Weeks Postpartum | 10.1016/j.neuroscience.2018.07.006 | Sweden | Europe and Central Asia |
| Luders | 2020 | From baby brain to mommy brain: Widespread gray matter gain after giving birth | https://doi.org/10.1016/j.cortex.2019.12.029 | Sweden | Europe and Central Asia |
| Luders | 2021 | Gray matter increases within subregions of the hippocampal complex after pregnancy | https://doi.org/10.1007/s11682-021-00463-2 | Sweden | Europe and Central Asia |
| Luders | 2021 | Postpartum Gray Matter Changes in the Auditory Cortex | https://doi.org/10.3390/jcm10235616 | Sweden | Europe and Central Asia |
| Luders | 2021 | Significant increases of the amygdala between immediate and late postpartum: Pronounced effects within the superficial subregion | 10.1002/jnr.24855 | Sweden | Europe and Central Asia |
| Luo | 2020 | Effects of normal pregnancy on maternal EEG, TCD, and cerebral cortical volume | https://doi.org/10.1016/j.bandc.2020.105526 | China | East Asia Pacific |
| Mangelsdorf | 2017 | Coping with Childbirth: Brain Structural Associations of Personal Growth Initiative | 10.3389/fpsyg.2017.01829 | Germany | Europe and Central Asia |
| Martinez-Garcia | 2021 | Do Pregnancy-Induced Brain Changes Reverse? The Brain of a Mother Six Years after Parturition | https://doi.org/10.3390/brainsci11020168 | Spain | Europe and Central Asia |
| Matsuda | 2011 | Processing of infant-directed speech by adults | 10.1016/j.neuroimage.2010.07.072 | Japan | East Asia Pacific |
| Matsuda | 2014 | Auditory observation of infant-directed speech by mothers: Experience-dependent interaction between language and emotion in the basal ganglia | 10.3389/fnhum.2014.00907 | Japan | East Asia Pacific |

| | | | | | |
|---|---|---|---|---|---|
| McEwen | 2021 | Glutamate levels in the medial prefrontal cortex of healthy pregnant women compared to non-pregnant controls | https://doi.org/10.1016/j.psyneuen.2021.105382 | Canada | North America |
| Montirosso | 2017 | Greater brain response to emotional expressions of their own children in mothers of preterm infants: an fMRI study | doi:10.1038/jp.2017.2 | Italy | Europe and Central Asia |
| Morgan | 2017 | Postpartum depressive symptoms moderate the link between mothers' neural response to positive faces in reward and social regions and observed caregiving | 10.1093/scan/nsx087 | USA | North America |
| Moser | 2013 | Limbic brain responses in mothers with post-traumatic stress disorder and comorbid dissociation to video clips of their children | https://doi.org/10.3109/10253890.2013.816280 | Switzerland | Europe and Central Asia |
| Moser | 2015 | BDNF methylation and maternal brain activity in a violence-related sample | 10.1371/journal.pone.0143427 | Switzerland | Europe and Central Asia |
| Moser | 2015 | The relation of general socio-emotional processing to parenting specific behavior: A study of mothers with and without posttraumatic stress disorder | 10.3389/fpsyg.2015.01575 | Switzerland | Europe and Central Asia |
| Moser | 2019 | Parental Reflective Functioning correlates to brain activation in response to video-stimuli of mother-child dyads: Links to maternal trauma history and PTSD | https://doi.org/10.1016/j.pscychresns.2019.09.005 | Switzerland | Europe and Central Asia |
| Moses-Kolko | 2010 | Abnormally reduced dorsomedial prefrontal cortical activity and effective connectivity with amygdala in response to negative emotional faces in postpartum depression | 10.1176/appi.ajp.2010.09081235 | USA | North America |
| Moses-Kolko | 2011 | Rapid habituation of ventral striatal response to reward receipt in postpartum depression | 10.1016/j.biopsych.2011.02.021 | USA | North America |
| Moses-Kolko | 2016 | The influence of motherhood on neural systems for reward processing in low income, minority, young women | 10.1016/j.psyneuen.2016.01.009 | USA | North America |
| Moses-Kolko | 2021 | Reduced postpartum hippocampal volume is associated with positive mother-infant caregiving behavior | 10.1016/j.jad.2020.12.014 | USA | North America |
| Musser | 2012 | The neural correlates of maternal sensitivity: an fMRI study | https://doi.org/10.1016/j.dcn.2012.04.003 | USA | North America |
| Nah | 2018 | Altered task-dependent functional connectivity patterns during subjective recollection experiences of episodic retrieval in postpartum women | https://doi.org/10.1016/j.nlm.2018.03.008 | South Korea | East Asia Pacific |
| Nah | 2018 | Data on subjective recollection effects reflected in large-scale functional connectivity patterns in postpartum women | https://doi.org/10.1016/j.nlm.2018.03.008 | South Korea | East Asia Pacific |
| Nitschke | 2004 | Orbitofrontal cortex tracks positive mood in mothers viewing pictures of their newborn infants | 10.1016/j.neuroimage.2003.10.005 | USA | North America |
| Noll | 2018 | Behavioral and neural correlates of parenting self-evaluation in mothers of young children | 10.1093/scan/nsy031 | USA | North America |
| Noriuchi | 2008 | The functional neuroanatomy of maternal love: mother's response to infant's attachment behaviors | 10.1016/j.biopsych.2007.05.018 | Japan | East Asia Pacific |
| Noriuchi | 2019 | The orbitofrontal cortex modulates parenting stress in the maternal brain | https://doi.org/10.1038/s41598-018-38402-9 | Japan | East Asia Pacific |
| Oatridge | 2002 | Change in brain size during and after pregnancy: study in healthy women and women with preeclampsia | DOI missing: link to paper https://www.ajnr.org/content/23/1/19.short | United Kingdom | Europe and Central Asia |
| O'Brien | 2021 | Is postnatal depression a distinct subtype of major depressive disorder? An exploratory study | https://doi.org/10.1007/s00737-020-01051-x | United Kingdom | Europe and Central Asia |
| Ojha | 2022 | Empathy for others versus for one's child: Associations with mothers' brain activation during a social cognitive task and with their toddlers' functioning | 10.1002/dev.22313 | USA | North America |

| | | | | | |
|---|---|---|---|---|---|
| Olsavsky | 2019 | Neural processing of infant and adult face emotion and maternal exposure to childhood maltreatment | doi: 10.1093/scan/nsz069 | USA | North America |
| Olsavsky | 2021 | Reported maternal childhood maltreatment experiences, amygdala activation and functional connectivity to infant cry | doi: 10.1093/scan/nsab005 | USA | North America |
| Orchard | 2023 | The maternal brain is more flexible and responsive at rest: effective connectivity of the parental caregiving network in postpartum mothers | https://doi.org/10.1038/s41598-023-31696-4 | Australia | East Asia Pacific |
| Ostrem | 2022 | Peripartum disease activity in moderately and severely disabled women with multiple sclerosis | 10.1177/20552173221104918 | USA, Canada, United Kingdom | Intercontinental |
| Parsons | 2017 | Duration of motherhood has incremental effects on mothers' neural processing of infant vocal cues: a neuroimaging study of women | 1038/s41598-017-01776-3 | Denmark | Europe and Central Asia |
| Raghunath | 2022 | Stronger brain activation for own baby but similar activation toward babies of own and different ethnicities in parents living in a multicultural environment | https://doi.org/10.1038/s41598-022-15289-1 | China | East Asia Pacific |
| Ranote | 2004 | The neural basis of maternal responsiveness to infants: an fMRI study | 10.1097/01.wnr.0000137078.64128.6a | United Kingdom | Europe and Central Asia |
| Rehbein | 2022 | Pregnancy and brain architecture: Associations with hormones, cognition and affect | 10.1111/jne.13066 | Germany | Europe and Central Asia |
| Rigo | 2019 | Brain Processes in Mothers and Nulliparous Women in Response to Cry in Different Situational Contexts: A Default Mode Network Study | https://doi.org/10.1080/15295192.2019.1555430 | Italy | Europe and Central Asia |
| Rodrigo | 2020 | Neglectful maternal caregiving involves altered brain volume in empathy-related areas | 10.1017/S0954579419001469 | Spain | Europe and Central Asia |
| Rupp | 2013 | Lower sexual interest in postpartum women: relationship to amygdala activation and intranasal oxytocin | 10.1016/j.yhbeh.2012.10.007 | USA | North America |
| Rupp | 2014 | Amygdala response to negative images in postpartum vs nulliparous women and intranasal oxytocin | 10.1093/scan/nss100 | USA | North America |
| Rutherford | 2003 | Magnetic resonance spectroscopy in pre-eclampsia: evidence of cerebral ischaemia | 10.1016/S1470-0328(03)00916-9 | United Kingdom | Europe and Central Asia |
| Rutherford | 2015 | Investigating Maternal Brain Structure and its Relationship to Substance Use and Motivational Systems | PMCID: PMC7974101 | USA | North America |
| Rutherford | 2019 | Gradient theories of brain activation: A novel application to studying the parental brain | https://doi.org/10.1007/s40473-019-00182-5 | USA | North America |
| Rutherford | 2020 | The Application of Connectome-Based Predictive Modeling to the Maternal Brain: Implications for Mother-Infant Bonding | 10.1093/cercor/bhz185 | USA | North America |
| Sambataro | 2021 | Altered dynamics of the prefrontal networks are associated with the risk for postpartum psychosis: a functional magnetic resonance imaging study | https://doi.org/10.1038/s41398-021-01351-5 | United Kingdom | Europe and Central Asia |
| Sasaki | 2020 | Cerebral diffusion kurtosis imaging to assess the pathophysiology of postpartum depression | https://doi.org/10.1038/s41598-020-72310-1 | Japan | East Asia Pacific |
| Schechter | 2012 | An fMRI study of the brain responses of traumatized mothers to viewing their toddlers during separation and play | doi:10.1093/scan/nsr069 | USA | North America |
| Schechter | 2015 | Methylation of NR3C1 is related to maternal PTSD, parenting stress and maternal medial prefrontal cortical activity in response to child separation among mothers with histories of violence exposure | 10.3389/fpsyg.2015.00690 | Switzerland | Europe and Central Asia |
| Schechter | 2017 | Maternal PTSD and corresponding neural activity mediate effects of child exposure to violence on child PTSD symptoms | https://doi.org/10.1371/journal.pone.0181066 | Switzerland | Europe and Central Asia |

| | | | | | |
|---|---|---|---|---|---|
| Schechter | 2017 | The association of serotonin receptor 3A methylation with maternal violence exposure, neural activity, and child aggression | https://doi.org/10.1016/j.bbr.2016.10.009 | Switzerland | Europe and Central Asia |
| Schnakenberg | 2021 | Examining early structural and functional brain alterations in postpartum depression through multimodal neuroimaging | https://doi.org/10.1038/s41598-021-92882-w | Germany | Europe and Central Asia |
| Schnakenberg | 2021 | The early postpartum period - Differences between women with and without a history of depression | https://doi.org/10.1016/j.jpsychires.2021.01.056 | Germany | Europe and Central Asia |
| Schneider | 2023 | Stress and reward in the maternal brain of mothers with borderline personality disorder: a script-based fMRI study | https://doi.org/10.1007/s00406-023-01634-6 | Germany | Europe and Central Asia |
| Seifritz | 2003 | Differential sex-independent amygdala response to infant crying and laughing in parents versus nonparents | 10.1016/S0006-3223(03)00697-8 | Switzerland | Europe and Central Asia |
| Shearrer | 2019 | The impact of elevated body mass on brain responses during appetitive prediction error in postpartum women | https://doi.org/10.1016/j.physbeh.2019.04.009 | USA | North America |
| Shimada | 2018 | Subclinical maternal depressive symptoms modulate right inferior frontal response to inferring affective mental states of adults but not of infants | https://doi.org/10.1016/j.jad.2017.12.031 | Japan | East Asia Pacific |
| Shimon-Raz | 2021 | Mother brain is wired for social moments | https://doi.org/10.7554/eLife.59436 | Israel | Middle East and North Africa |
| Shin | 2018 | Disturbed retrieval network and prospective memory decline in postpartum women | 10.1038/s41598-018-23875-5 | South Korea | East Asia Pacific |
| Silver | 2018 | White matter integrity in medication-free women with peripartum depression: a tract-based spatial statistics study | https://doi.org/10.1038/s41386-018-0023-y | USA | North America |
| Silverman | 2007 | Neural dysfunction in postpartum depression: an fMRI pilot study | https://doi.org/10.1017/S1092852900015595 | USA | North America |
| Silverman | 2011 | The neural processing of negative emotion postpartum: a preliminary study of amygdala function in postpartum depression | 10.1007/s00737-011-0226-2 | USA | North America |
| Stickel | 2019 | Cumulative cortisol exposure in the third trimester correlates with postpartum mothers' neural response to emotional interference | https://doi.org/10.1016/j.biopsycho.2019.02.008 | Germany | Europe and Central Asia |
| Strathearn | 2008 | What's in a smile? Maternal brain responses to infant facial cues | www.pediatrics.org/cgi/doi/10.1542/peds.2007-1566 | USA | North America |
| Strathearn | 2009 | Adult attachment predicts maternal brain and oxytocin response to infant cues | 10.1038/npp.2009.103 | USA | North America |
| Strathearn | 2013 | Mothers' amygdala response to positive or negative infant affect is modulated by personal relevance | 10.3389/fnins.2013.00176 | USA | North America |
| Sui | 2023 | Decreased gray matter volume in the right middle temporal gyrus associated with cognitive dysfunction in preeclampsia superimposed on chronic hypertension | 10.3389/fnins.2023.1138952 | China | East Asia Pacific |
| Swain | 2008 | Maternal brain response to own baby-cry is affected by cesarean section delivery | 10.1111/j.1469-7610.2008.01963.x. | USA | North America |
| Swain | 2017 | Parent-child intervention decreases stress and increases maternal brain activity and connectivity during own baby-cry: An exploratory study | doi:10.1017/S0954579417000165. | USA | North America |
| Swain | 2019 | Early postpartum resting-state functional connectivity for mothers receiving buprenorphine treatment for opioid use disorder: A pilot study | doi:10.1111/jne.12770 | USA | North America |
| Swain | 2021 | Reduced Child-Oriented Face Mirroring Brain Responses in Mothers With Opioid Use Disorder: An Exploratory Study | 10.3389/fpsyg.2021.770093 | USA | North America |

| | | | | | |
|---|---|---|---|---|---|
| Swain | 2023 | Brain circuits for maternal sensitivity and pain involving anterior cingulate cortex among mothers receiving buprenorphine treatment for opioid use disorder | 10.1111/jne.13316 | USA | North America |
| Uher | 2022 | Pregnancy-induced brain magnetic resonance imaging changes in women with multiple sclerosis | 10.1111/ene.15245 | Czech Republic | Europe and Central Asia |
| Wan | 2014 | The neural basis of maternal bonding | https://doi.org/10.1371/journal.pone.0088436 | United Kingdom | Europe and Central Asia |
| Wonch | 2016 | Postpartum depression and brain response to infants: Differential amygdala response and connectivity | http://dx.doi.org/10.1080/17470919.2015.1131193 | Canada | North America |
| Wright | 2017 | Mothers Who Were Neglected in Childhood Show Differences in Neural Response to Their Infant's Cry | 10.1177/1077559516683503 | USA | North America |
| Xiao-Juan | 2011 | Increased Posterior Cingulate, Medial Frontal and Decreased Temporal Regional Homogeneity in Depressed Mothers. A Resting-State Functional Magnetic Resonance Study | 10.1016/j.proenv.2011.10.112 | China | East Asia Pacific |
| Xu | 2023 | Consistent functional abnormalities in patients with postpartum depression | https://doi.org/10.1016/j.bbr.2023.114467 | China | East Asia Pacific |
| Yang | 2022 | Oxygen extraction fraction (OEF) assesses cerebral oxygen metabolism of deep gray matter in patients with pre-eclampsia | https://doi.org/10.1007/s00330-022-08713-7 | China | East Asia Pacific |
| Yang | 2023 | Cortical and subcortical morphological alterations in postpartum depression | https://doi.org/10.1016/j.bbr.2023.114414 | China | East Asia Pacific |
| Zhang | 2019 | Brain Structural Plasticity Associated with Maternal Caregiving in Mothers: A Voxel- and Surface-Based Morphometry Study | 10.1159/000506258 | China | East Asia Pacific |
| Zhang | 2020 | Aberrant resting-state interhemispheric functional connectivity in patients with postpartum depression | https://doi.org/10.1016/j.bbr.2020.112483 | China | East Asia Pacific |
| Zhang | 2020 | Brain Responses to Emotional Infant Faces in New Mothers and Nulliparous Women | https://doi.org/10.1038/s41598-020-66511-x | China | East Asia Pacific |
| Zhang | 2022 | Abnormal Voxel-Based Degree Centrality in Patients With Postpartum Depression: A Resting-State Functional Magnetic Resonance Imaging Study | 10.3389/fnins.2022.914894 | China | East Asia Pacific |
| Zhang | 2022 | Decreased brain functional connectivity associated with cognitive dysfunction in women with second pregnancy | 10.3389/fnagi.2022.963943 | China | East Asia Pacific |
| Zhang | 2022 | Improved Interhemispheric Functional Connectivity in Postpartum Depression Disorder: Associations With Individual Target-Transcranial Magnetic Stimulation Treatment Effects | 10.3389/fpsyt.2022.859453 | China | East Asia Pacific |
| Zhang | 2023 | Assessing Cerebral Oxygen Metabolism Changes in Patients With Preeclampsia Using Voxel-Based Morphometry of Oxygen Extraction Fraction Maps in Magnetic Resonance Imaging | 10.3348/kjr.2022.0652 | China | East Asia Pacific |
| Zheng | 2018 | Disrupted Spontaneous Neural Activity Related to Cognitive Impairment in Postpartum Women | 10.3389/fpsyg.2018.00624 | China | East Asia Pacific |
| Zheng | 2020 | Disruption within brain default mode network in postpartum women without depression | http://dx.doi.org/10.1097/MD.0000000000020045 | China | East Asia Pacific |
